# Fetal influence on maternal pregnancy through the *Peg3* imprinted domain

**DOI:** 10.64898/2025.11.28.691217

**Authors:** Joomyeong Kim

## Abstract

Genomic imprinting plays critical roles in the reproduction of mammalian species. In this study, potential roles of the *Peg3* imprinted domain in mammalian reproduction were analyzed with the mutant mouse model lacking the oocyte-specific alternative promoter U1, which exhibits the abnormal biallelic expression or 2x gene dosage of *Peg3*. According to the results, the pregnant females displayed several mutant phenotypes related to reproduction, including delayed parturition, and defects in placentophagy and nest-building behaviors, causing subsequent neonatal lethality. These phenotypes were observed from the homozygote females delivering the entire litter with 2x *Peg3*, but not from the heterozygotes producing a half of litter with 2x *Peg3*, suggesting the presence of maximal tolerable gene dosage of *Peg3* for healthy pregnancy as well as potential fetal influence on maternal pregnancy through the gene dosage of the *Peg3* domain. Detailed analyses also revealed partial embryonic lethality among the female with 2x *Peg3*, suggesting the susceptibility of the female to the elevated gene dosage of *Peg3*. Overall, this highlights that the gene dosage of the *Peg3* domain is critical for the successful mammalian reproduction.

## Introduction

In mammalian genomes, a small subset of autosomal genes—about 200—is expressed predominantly from one allele due to the epigenetic mechanism known as genomic imprinting, in which one allele is repressed through DNA methylation and histone modifications [1,2]. This mechanism is found mainly in eutherian mammals and is believed to have co-evolved with their distinctive reproductive strategy involving placentation and viviparity [3-5]. Consistent with this idea, many imprinted genes are highly expressed in tissues important for mammalian reproduction, particularly in the placenta [1,2,6]. Studies have shown that the optimal gene dosage of several placental imprinted genes, including *Phlda2* and *Igf2,* is critical for maternal care behaviors and the physiological state of pregnant females, suggesting potential roles for imprinted genes in coordinating communication between the fetus and the mother during pregnancy [7,8]. Nevertheless, the majority of imprinted genes remain poorly studied with respect to their reproductive roles, even though the unusual reproductive scheme of mammals may have been a major driving force for the emergence of genomic imprinting in this lineage [3-5].

*Peg3* (Paternally expressed gene 3) is the founding member of an evolutionarily conserved 500-kb imprinted domain located on human chromosome 19q13.4 and proximal mouse chromosome 7 [9]. This imprinted domain contains the paternally expressed genes *Peg3*, *Usp29*, *Zfp264*, and *APeg3*, and the maternally expressed genes *Zim1*, *Zim2*, and *Zim3* [10]. Although the overall genomic structure of the domain has been well preserved in mammals, *Peg3* is the only gene that has maintained its open reading frame (ORF) throughout mammalian evolution, suggesting that its *in vivo* functions are particularly critical for species survival [10]. Imprinting of this domain is regulated by two *cis*-regulatory elements: the Peg3-DMR (Differentially Methylated Region) and the oocyte-specific alternative promoter U1. The Peg3-DMR, which encompasses the bidirectional promoter for *Peg3* and *Usp29*, functions as the imprinting control region (ICR) for the domain; its deletion disrupts the monoallelic expression of all genes within the domain [11,12]. This ICR normally acquires gametic DNA methylation during oogenesis through transcription-mediated mechanisms involving the U1 promoter, and deletion of this promoter results in biallelic expression or 2x dosage of *Peg3* [13].

The potential roles of *Peg3* in reproduction have been explored using mutant mouse models and a limited number of human studies. In mice, loss-of-function mutations that eliminate *Peg3* expression with 0x dosage typically lead to reduced growth rates and defects in maternal care behaviors in pregnant females [14-16]. Follow-up studies have shown that these behavioral defects are likely caused by disruptions in the oxytocin circuitry, which appears to be directly regulated by PEG3, a DNA-binding transcriptional repressor [17]. In humans, *PEG3* expression levels in the placenta have been associated with prenatal depression in pregnant women, suggesting a potential link between fetal *PEG3* dosage and maternal mental health [18]. As part of ongoing efforts, the present study aimed to investigate the effects of biallelic expression or 2x dosage of *Peg3* on reproduction using the mutant mouse strain lacking the U1 promoter [13]. The results show that homozygous females for the U1 deletion exhibit several reproduction-related phenotypes, including delayed parturition and defects in placentophagy and nest-building behaviors. Furthermore, these defects appear to originate from the fetus carrying 2x *Peg3*, rather than from the pregnant female herself, suggesting potential fetal influence on maternal pregnancy through the *Peg3* imprinted domain.

## Results

### Gene dosage of the *Peg3* domain controlled by two *cis*-regulatory elements

The mouse *Peg3* imprinted domain contains four paternally expressed and three maternally expressed genes (**Figure 1A**) [10]. The imprinting, or monoallelic expression, of this domain is regulated by two *cis*-regulatory elements: the Peg3-DMR (Differentially Methylated Region), which flanks the bidirectional promoter for paternally expressed *Peg3* and *Usp29*, and the oocyte-specific alternative promoter U1 for *Peg3*. The Peg3-DMR functions as the ICR (Imprinting Control Region), and paternal deletion of this ICR—referred to as the KO2 allele—results in the absence of expression or 0x dosage of both *Peg3* and *Usp29*, along with de-repression of the paternal allele of *Zim1*, causing its biallelic expression or 2x gene dosage (**Figure 1C**) [11,12]. Targeting of DNA methylation to the Peg3-DMR is thought to be mediated by U1-driven transcription during oogenesis. Accordingly, maternal deletion of this alternative promoter—referred to as the U1 allele—results in a lack of methylation at the Peg3-DMR, which in turn leads to reactivation of the maternal allele and biallelic expression or 2x gene dosage of *Peg3* and *Usp29* (**Figure 1B**) [13]. However, this U1-dependent methylation mechanism is not fully efficient; approximately 10–20% of mature oocytes and their progeny produced by homozygous U1 females retain normal DNA methylation at the Peg3-DMR and thus maintain the normal 1x gene dosage of the *Peg3* domain [13]. Because the U1 promoter is active only during oogenesis, paternal transmission of the U1 allele has no effect on the maternal-specific DNA methylation of the Peg3-DMR or on the overall gene dosage of the *Peg3* domain [19]. Therefore, both heterozygous and homozygous carriers of the U1 allele exhibit the same *Peg3* gene dosage, regardless of the genotype of the paternal allele (**Figure 1B**).

**Figure 1.**
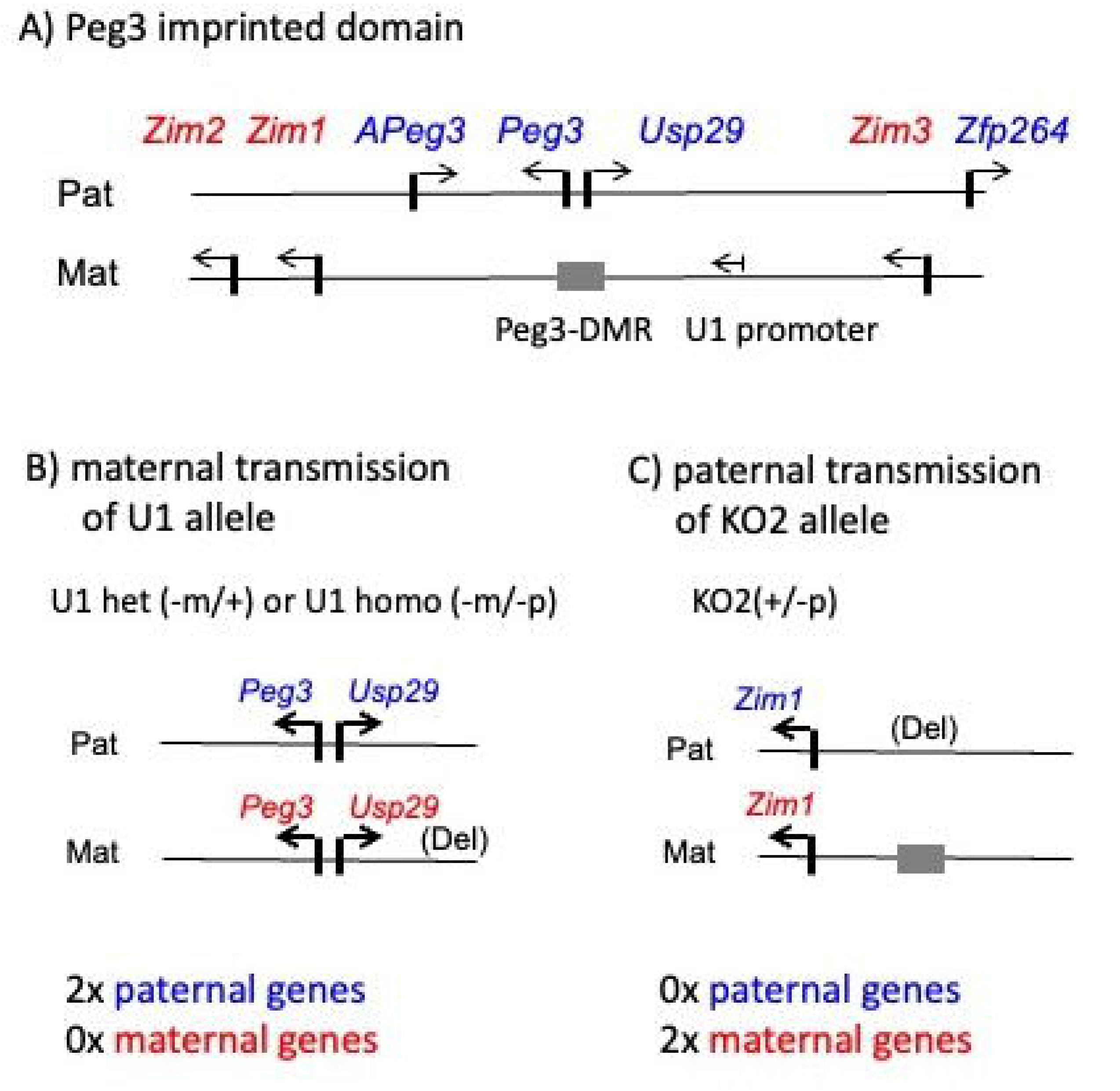
Mouse *Peg3* imprinted domain and two mutant alleles. (**A**) The *Peg3* imprinted domain contains seven genes, indicated by arrows. Paternally expressed genes (*Peg3, Usp29, Zfp264, APeg3*) are shown in blue, and maternally expressed genes (*Zim1, Zim2, Zim3*) are shown in red. The 4-kb Peg3-DMR, shown as a gray box, controls imprinting for the entire domain. The oocyte-specific alternative promoter U1, located 20 kb upstream of the Peg3-DMR, directs *de novo* DNA methylation to the maternal allele during oogenesis and is indicated with a small arrow. (**B**) Maternal transmission of the U1 allele (U1 deletion) reactivates *Peg3* and *Usp29* from the maternal allele (2x dosage) and causes down-regulation of the maternally expressed *Zim1* (0x dosage). (**C**) Paternal transmission of the KO2 allele (Peg3-DMR deletion) abolishes transcription of *Peg3* and *Usp29* (0x dosage) and reactivates the paternal *Zim1* allele (2x dosage).

### Breeding schemes for potential gene dosage effects of the *Peg3* domain

Potential effects of different gene dosages of the *Peg3* domain on reproduction were analyzed using six breeding schemes that employed two mutant strains carrying the U1 and KO2 alleles (**Figure 2**). As described earlier, maternal transmission of the U1 allele and paternal transmission of the KO2 allele change the gene dosage of the *Peg3* domain. Accordingly, female heterozygotes and homozygotes for the U1 allele were individually crossed with three types of males: U1 homozygotes (−/−), wild type (+/+), and KO2 heterozygotes (−/+). In the case of the KO2 allele, male homozygotes are barely viable and therefore not available for breeding experiments [12]. The expected gene dosages of potential litters from the first three breeding schemes (Schemes 1–3), which involve U1 heterozygous females, are as follows. Half of the progeny from U1 heterozygous females are expected to inherit the U1 allele and thus potentially have the 2x dosage of *Peg3*. However, because approximately 10–20% of eggs and resulting progeny retain normal DNA methylation on the Peg3-DMR, they maintain the normal 1x gene dosage. This results in the overall expected dosage range of 1.0–1.5x for Schemes 1 and 2. In contrast, male heterozygotes for the KO2 allele are expected to produce litters in which half of the offspring have the 0x dosage of *Peg3*, yielding the overall dosage range of 0.5–1.0x for Scheme 3. The breeding schemes involving U1 homozygous females (Schemes 4–6) are expected to produce similar dosage ranges, except with a 0.5x increase compared with the U1 heterozygous schemes. This is because the entire litter—not just half—will inherit the U1 allele and therefore potentially have the 2x dosage of *Peg3*. It is also important to note that the gene dosage of the females themselves can be either 1x or 2x in these breeding experiments, even though both heterozygotes and homozygotes carry the maternally transmitted U1 allele (**Figure 2**).

**Figure 2.**
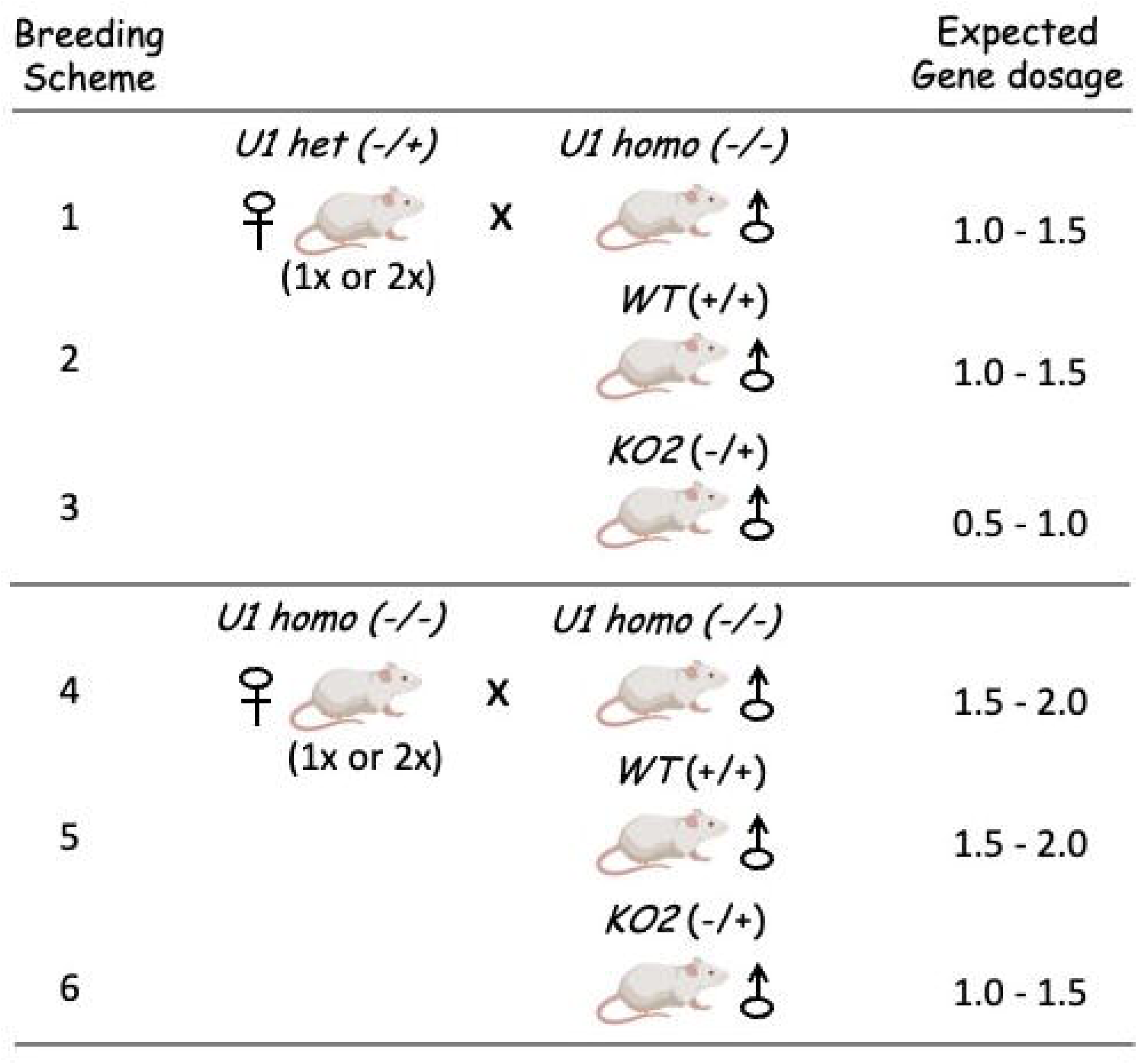
Breeding schemes generating variable *Peg3* gene dosages. Female heterozygotes (U1 het; Schemes 1–3) and homozygotes (U1 homo; Schemes 4–6) for the U1 allele were crossed with three types of males: U1 homozygotes (Schemes 1 and 4), wild-type littermates (Schemes 2 and 5), and KO2 heterozygotes (Schemes 3 and 6). These breeding combinations generate litters with distinct *Peg3* gene dosage ranges, summarized in the final column.

### Breeding results for potential gene dosage effects of the *Peg3* domain

The litters produced from the six breeding schemes were analyzed for their litter size, lethality during the first week after birth, sex ratio, and average gene dosage of *Peg3* (**Table 1**). The average gene dosage per pup was measured using DNA isolated from each individual pup with the restriction-enzyme–based protocol COBRA (Combined Bisulfite and Restriction Analysis) [20]. The first three breeding schemes, which involved approximately 40 female U1 heterozygotes, were derived from archived records and DNA samples collected in two previous studies [13,21]. The remaining three schemes were generated in the current study using 75 female U1 homozygotes. Within each female group, about one-third of the females exhibited 1x gene dosage and the remaining two-thirds exhibited 2x gene dosage. Females of both dosage categories were randomly selected for each breeding experiment. The results from the six breeding schemes were summarized, leading to the following conclusions. First, the actual gene dosages observed in each scheme fell within their expected ranges. The sex ratios were also normal, showing an overall 1:1 ratio between males and females, although three schemes (Schemes 1, 2, and 6) showed slightly more males than females. These findings suggest that embryogenesis does not exhibit any obvious bias or selection toward specific gene dosages or sexes (**Table 1**). In contrast, litter sizes varied among schemes: four schemes (Schemes 2, 4, 5, and 6) yielded approximately six pups per litter, whereas the remaining two (Schemes 1 and 3) produced 8–9 pups per litter. The litter sizes of Schemes 1 and 3 are consistent with the normal range for the C57BL/6J background of the U1 mutant strain [22]. The reduced litter sizes of the other four schemes—smaller by two to three pups—suggest potential embryonic lethality. Notably, the schemes with normal litter sizes (Schemes 1 and 3) also displayed gene dosages within the normal range (1.11x and 0.86x, respectively), whereas the schemes with reduced litter sizes (Schemes 2, 4, 5, and 6) showed higher gene dosages (1.30x, 1.69x, 1.70x, and 1.27x, respectively). This correlation suggests that elevated gene dosage of *Peg3* may contribute to embryonic lethality, a possibility addressed further below.

**Table 1.**
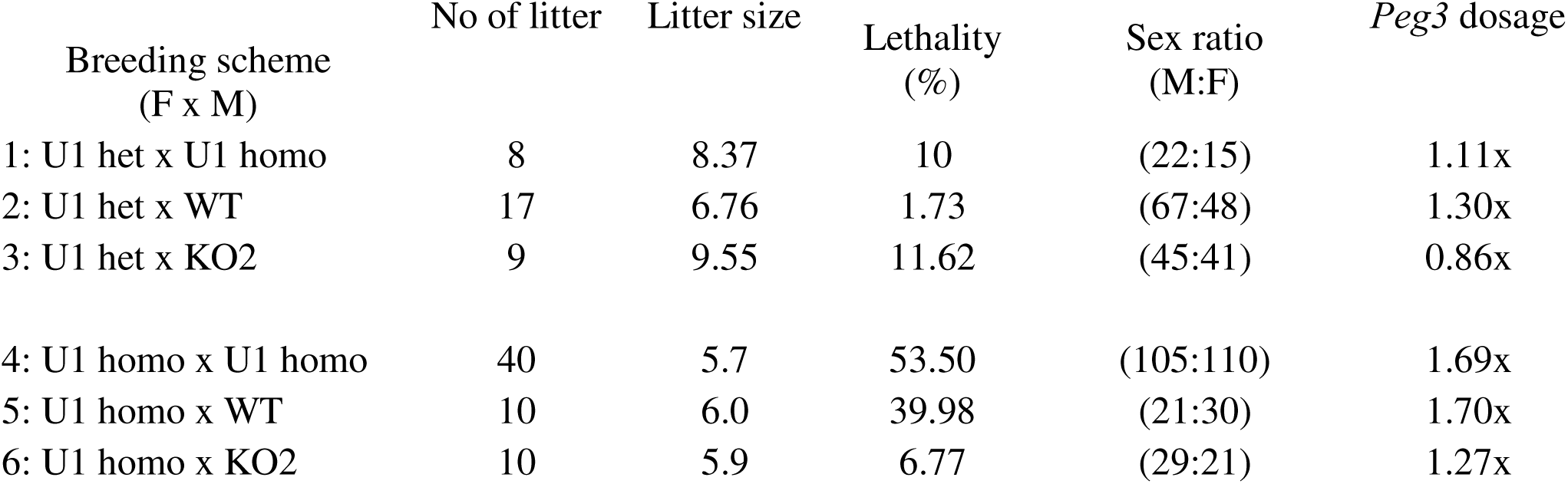
Breeding results.

Second, two breeding schemes—Schemes 4 and 5—displayed unusually high levels of neonatal lethality, 53.50% and 39.98%, respectively, compared with only 1.73% to 11.62% in the other schemes. These two schemes also exhibited substantially higher gene dosages among their pups (1.69x and 1.70x) relative to the others (ranging from 0.86x to 1.30x). Anecdotally, similar high neonatal lethality has frequently been observed during routine colony maintenance of this mutant line: pups from U1 homozygous females often die within a couple of days after birth for unknown reasons. These pups are often found scattered in the cage with empty stomachs, suggesting that maternal care defects may contribute to neonatal lethality. Indeed, litters from U1 homozygous females are often rescued by fostering to unrelated nursing females. Importantly, such high lethality has not been observed in other breeding schemes, especially those involving U1 heterozygous females, even though both heterozygous and homozygous females carry the same maternally transmitted U1 allele [13,21]. This pattern is consistent with the low neonatal lethality observed in Schemes 1–3. Thus, the data from this study resolve a long-standing question: the high neonatal lethality is closely associated with the gene dosage of the pups, rather than with the gene dosage of the pregnant females. This conclusion is further supported by the low lethality observed in Scheme 6 (6.77%), in which pups exhibited only a moderate gene dosage (1.27x), despite being carried by U1 homozygous females. Overall, these results indicate that the pups’ *Peg3* gene dosage—rather than that of the dams—is strongly associated with, and potentially responsible for, neonatal lethality, possibly by influencing pregnancy-related behaviors or physiological functions of the mothers.

### Mutant phenotypes contributing to neonatal lethality

The observed neonatal lethality was further examined by analyzing several mutant phenotypes frequently displayed by female U1 homozygotes. These included delayed parturition, defects in placentophagy (placenta-eating), and abnormalities in nest-building behavior—traits known to be critical for the survival of newborn pups [24-27]. Each phenotype was carefully monitored during the breeding experiments of Schemes 4–6, and archived records from Schemes 1–3 were retrospectively analyzed to detect the same phenotypes [13,21]. The frequency of each phenotype in each breeding scheme was then correlated with the gene dosage of both the pregnant females and their pups.

Delayed parturition: delayed parturition was analyzed by measuring the gestation length of each breeding pair in Schemes 4–6 (**Table 2**). A total of 40 U1 homozygous females were followed across two pregnancies, yielding 72 gestation-length measurements. In mice with a C57BL/6J background, a gestation length of 19.5 days is considered normal, with lengths beyond this threshold classified as abnormal and indicative of potential parturition defects [23].

**Table 2.**
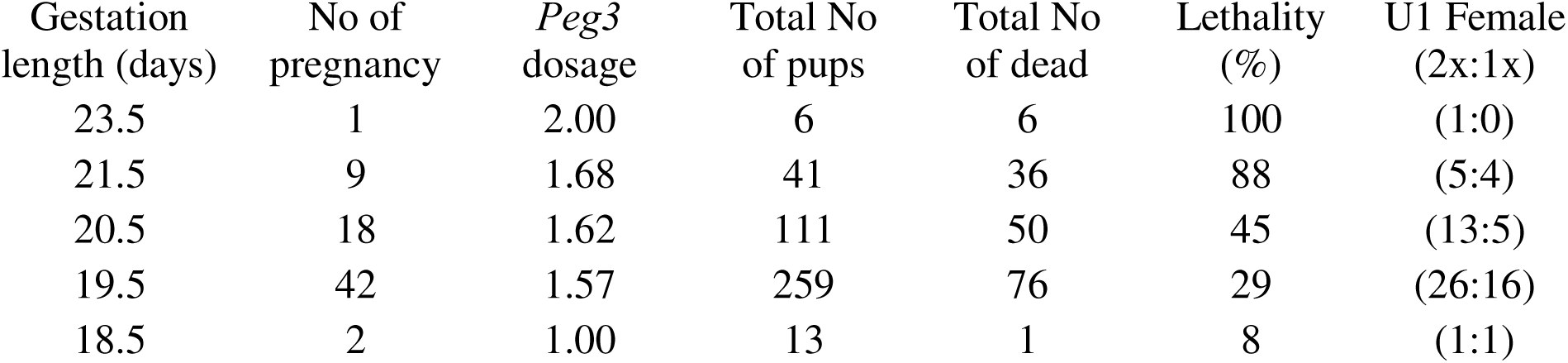
Delayed parturition among U1 homozygous females.

Approximately two-thirds of pregnancies (42/72) exhibited normal gestation (19.5 days), while the remaining one-third (28/72) were delayed, ranging from 20.5 to 23.5 days. Normal pregnancies displayed 29% neonatal lethality, whereas delayed pregnancies exhibited dramatically elevated lethality, ranging from 45% to 100%, indicating that delayed parturition is a major contributor to neonatal loss. In terms of gene dosage, no difference was detected between pregnant females in the normal and delayed groups: both 1x and 2x females were evenly represented across gestational categories (**Table 2**). By contrast, the average gene dosage of pups was higher in the delayed categories (1.62x–2.0x for 20.5–23.5 days) relative to the normal category (1.57x for 19.5 days). This trend is further supported by archived data from Schemes 1–3, which involved female U1 heterozygotes and showed no evidence of delayed parturition, consistent with the lower lethality observed in those litters (**Table 1**) [13,21]. Overall, delayed parturition appears to be a primary defect driving the high neonatal lethality in Schemes 4 and 5, and this phenotype is closely associated with increased gene dosage in the pups rather than in the pregnant females.

Defective placentophagy: female U1 homozygotes also frequently exhibited defects in placentophagy, a conserved maternal behavior essential for pup survival in placental mammals [26,27]. In cases of defective placentophagy, pups were found scattered in the cage with placentas still attached (**Figure 3A**). When placentophagy occurred normally, pups were fully cleaned and placed within a properly constructed nest (**Figure 3B**). Among 40 monitored U1 homozygous females, 18 displayed defective placentophagy during either their first or second pregnancy. This defect was more frequently observed among females with delayed parturition: 12 of the 18 displaying defective placentophagy also had gestations exceeding 19.5 days. Litters from females with defective placentophagy tended to show higher neonatal lethality and elevated pup gene dosage compared to those from females with normal placentophagy (**Figure 3**, lower panels). However, the defect occurred at similar frequencies among 1x and 2x females, indicating that defective placentophagy is also associated with increased pup gene dosage, rather than maternal dosage.

**Figure 3.**
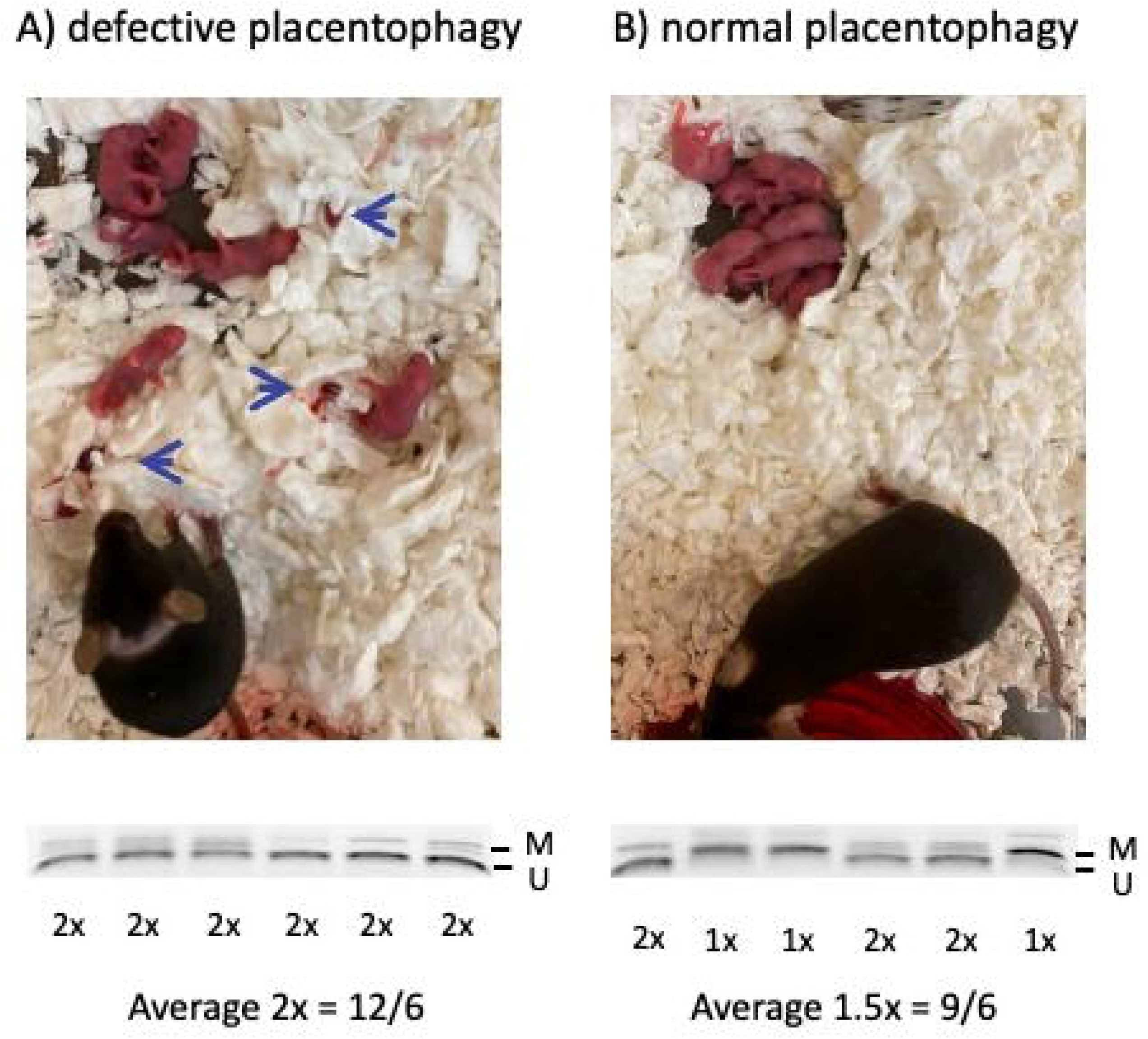
Defective placentophagy in U1 homozygous females. (**A**) Representative litter from a U1 homozygous female displaying defective placentophagy. Pups were dispersed throughout the cage, and uneaten placentas were attached to bedding (blue arrows; top). COBRA analysis showed unmethylated Peg3-DMR alleles, indicating that all pups carried 2x *Peg3* (bottom). (**B**) Representative litter from a U1 homozygous female with normal placentophagy. Pups were cleaned and placed within a well-formed nest (top). Gene dosage analysis showed that half of the pups carried 2x *Peg3* and the remaining half carried 1x *Peg3* (bottom), resulting in an average dosage of 1.5x *Peg3*.

Nest-building behavior: two distinct nest-building strategies were observed among U1 homozygous females (**Figure 4**). One group constructed nests underneath the plastic compartment holding food and water, placing the nests farther inside the cage, a location predicted to offer greater protection for newborn pups (**Figure 4A–B**). In contrast, another group failed to build their own nests prior to parturition and instead relied on the igloo-shaped plastic shelter provided at the front of the cage (**Figure 4C–D**). Because females must frequently leave this structure to access food and water, this nesting strategy was predicted to be less favorable for pup survival. Supporting this idea, more than 80% of nursing females unrelated to this study built nests beneath the plastic compartment. Survey results, interestingly, demonstrated that nest-building behavior strongly correlated with maternal gene dosage. Most U1 females with 1x dosage built their own nests (19 of 25), whereas nearly all females with 2x dosage failed to do so (37 of 39). However, nest-building behavior did not correlate with pup gene dosage, nor were differences in neonatal lethality detected between the two behavioral categories. Thus, contrary to initial predictions, nest construction did not significantly contribute to neonatal loss.

**Figure 4.**
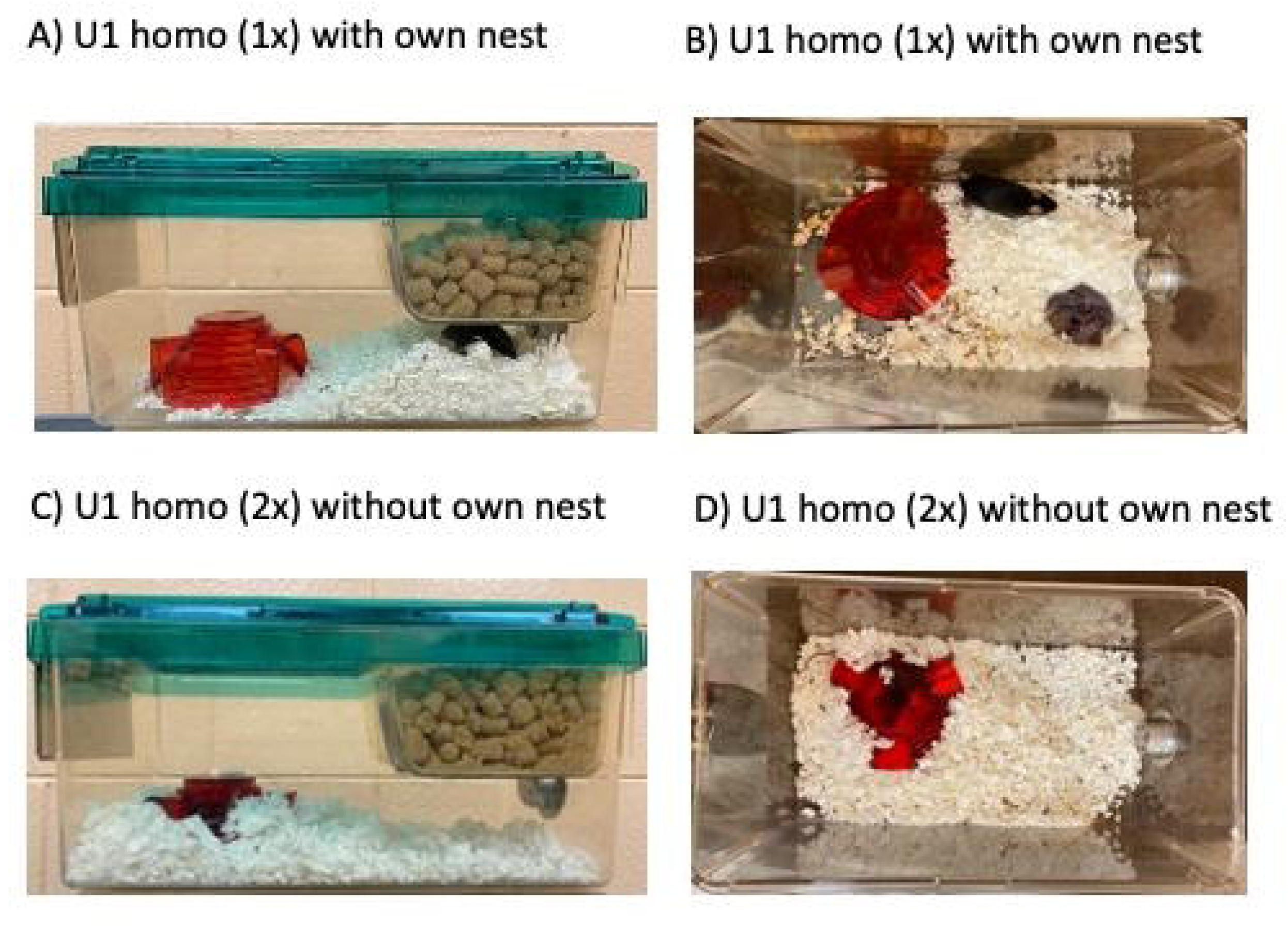
Defective nest-building behavior in U1 homozygous females. (**A–B**) Most U1 homozygous females with 1x *Peg3* built nests underneath the plastic food-and-water chamber, separate from the provided red igloo-shaped shelter. (**C–D**) In contrast, U1 homozygous females with 2x *Peg3* generally failed to construct nests at parturition and instead relied on the igloo-shaped plastic shelter originally provided during breeding setup.

In summary, these phenotypic analyses indicate that delayed parturition and defective placentophagy are the primary defects contributing to neonatal lethality, and both are closely associated with increased *Peg3* gene dosage in the pups. In contrast, while nest-building behavior correlates strongly with the gene dosage of pregnant females, it does not appear to influence neonatal survival.

### Embryonic lethality associated with U1 females with 2x gene dosage

Breeding results from Schemes 4–6 (**Table 1**) indicated that female U1 homozygotes consistently produced smaller litters, averaging 6 pups per litter—2 to 3 fewer than the expected litter size. This pattern suggests the presence of embryonic lethality. To investigate this possibility, a series of timed-mating experiments were performed using 25 U1 homozygous females. These matings generated 11 litters of embryos collected at developmental stages ranging from 12.5 to 17.5 days post-coitum (dpc). Among the 95 embryos harvested from these litters, 25 were undergoing resorption, corresponding to an average of approximately 2 resorbing embryos per litter (25/11). This observation is consistent with the earlier prediction that 2–3 embryos per litter fail to develop successfully. Because resorbing embryos were recovered, the lethality likely occurred during the organogenesis stage (9.5–14.5 dpc), as embryonic losses before this stage typically cannot be detected as identifiable resorbed tissues. A representative litter containing four dead and five live embryos is shown in **Figure 5**. DNA extracted from embryos was used to determine sex and *Peg3* gene dosage (**Figure 5C**). All 11 embryo litters were analyzed similarly, and findings were summarized by categorizing embryos according to sex and *Peg3* gene dosage (**Figure 5D**). Across all litters, the total numbers of female and male embryos were 53 and 42, respectively, approximating the expected 1:1 sex ratio. Additionally, 67 embryos carried 2x gene dosage, whereas 28 carried 1x dosage—frequencies consistent with those typically observed in U1 homozygote matings [13,21]. Notably, among the 25 dead embryos, the majority were females with 2x gene dosage: 18 of the 25 resorbing embryos belonged to this category. This pattern is striking and unexpected, given that the neonatal data from Schemes 4–6 showed no evidence of sex ratio distortion (**Table 1**). Nevertheless, these results strongly suggest that female embryos may be particularly susceptible to the elevated *Peg3* gene dosage, leading to increased vulnerability during embryogenesis.

**Figure 5.**
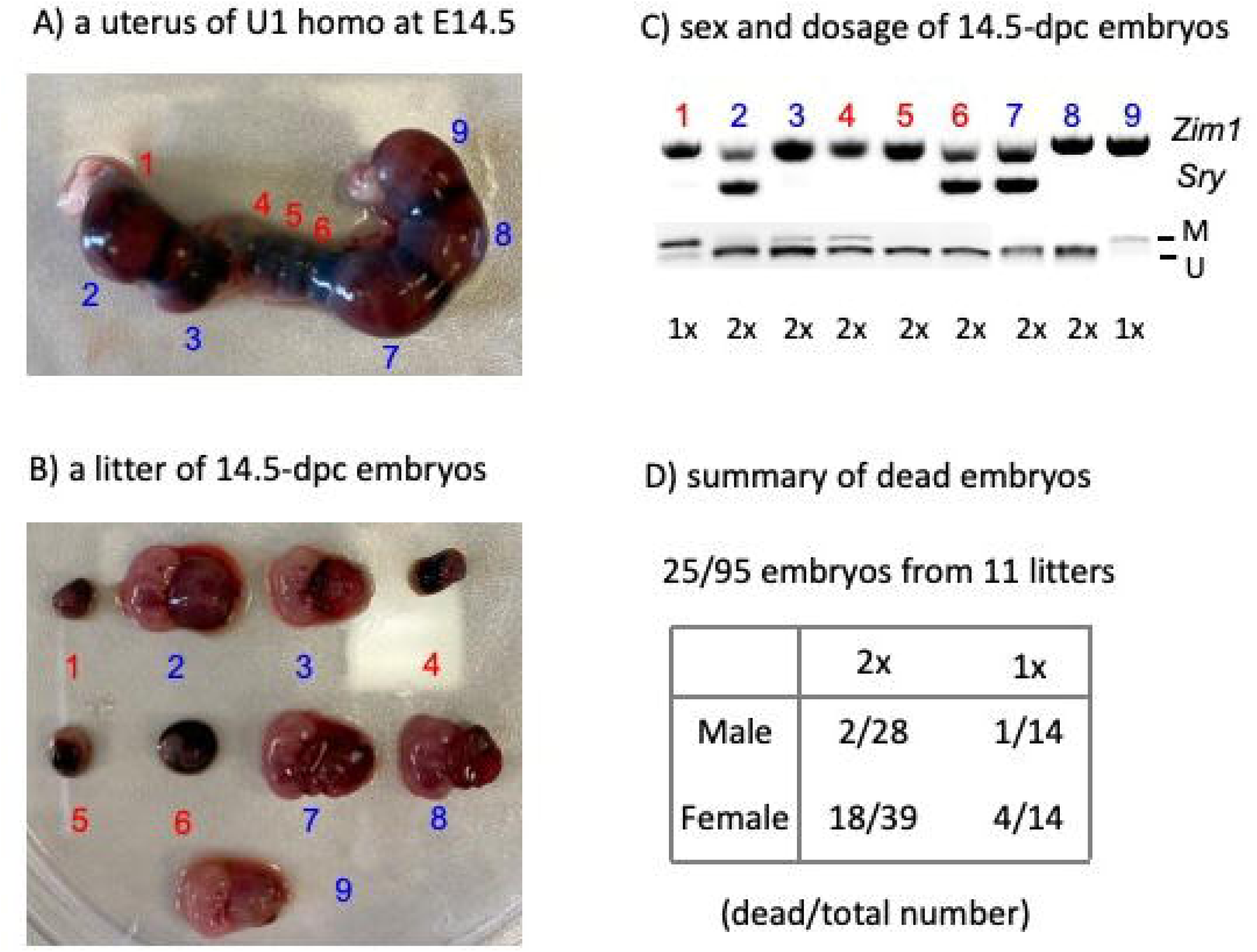
Female-specific lethality among U1 homozygous embryos. (**A–B**) Uterus collected from a U1 homozygous female bred with a U1 homozygous male. Nine embryos were recovered: five viable (blue) and four dead (red). (**C**) DNA from each embryo was used for sex determination *via Sry* PCR with *Zim1* as an autosomal control. *Peg3* gene dosage was assessed by COBRA, revealing methylated versus unmethylated Peg3-DMR alleles. (**D**) Summary table of sex and *Peg3* gene dosage for 95 embryos obtained from 11 timed-mating litters involving U1 homozygous females and males.

## Discussion

In the current study, we investigated the gene-dosage effects of the *Peg3* imprinted domain on mammalian reproduction using six breeding schemes involving two mutant mouse strains, the U1 and KO2 lines. The results support several major conclusions. First, two pregnancy-related phenotypes—delayed parturition and defective placentophagy—were the primary contributors to the neonatal lethality observed in litters produced by U1 homozygous females. Importantly, these phenotypes were strongly associated with the gene dosage of the pups, rather than that of the pregnant females. In contrast, nest-building behavior correlated with the gene dosage of the females but did not substantially contribute to neonatal lethality. Second, timed-mating experiments revealed that female embryos were particularly susceptible to elevated *Peg3* dosage, indicating a sex-specific sensitivity during embryogenesis. Collectively, these findings highlight that the gene dosage of the *Peg3* imprinted domain plays a critical role in mammalian reproduction, especially through its effects on female physiology.

The two major phenotypes—delayed parturition and defective placentophagy—clearly underlie the high levels of neonatal lethality in U1 homozygous matings (**Table 1**, **Table 2**, **Figure 3**, **Figure 4**). Surprisingly, the gene dosage of the pups, rather than that of their mothers, was the dominant factor associated with these defects. This unexpected outcome suggests that fetal genotype can influence maternal reproductive physiology through the *Peg3* imprinted domain (**Figure 6A**). This inference is consistent with the known biology of mammalian reproduction: females are primed for pregnancy-associated activities such as parturition, placentophagy, nest-building, nursing, and milk production, and this priming requires precise communication between the fetus and the mother [24-27]. Such communication is mediated by the placenta, which produces multiple hormones and signaling molecules essential for maintaining pregnancy [28,29]. Several genes within the *Peg3* imprinted domain, including *Peg3* and *Zim1*, are highly expressed in the placenta [30], and both gene products function as DNA-binding transcriptional regulators. Notably, PEG3 has been shown to bind promoters of numerous genes, including those involved in controlling progesterone levels—a hormone central to pregnancy maintenance and parturition [31,32]. Thus, elevated *Peg3* dosage may disrupt the expression of genes required for proper maternal priming, leading to failures in parturition timing and placentophagy (**Figure 6B**). Comparable fetal–maternal communication defects have been documented in other imprinted genes, such as *Igf2* and *Phlda2*, further supporting this model [7,8]. Taken together, these observations reinforce the hypothesis that genomic imprinting may have evolved, in part, to regulate critical aspects of fetal–maternal communication during pregnancy [33].

**Figure 6.**
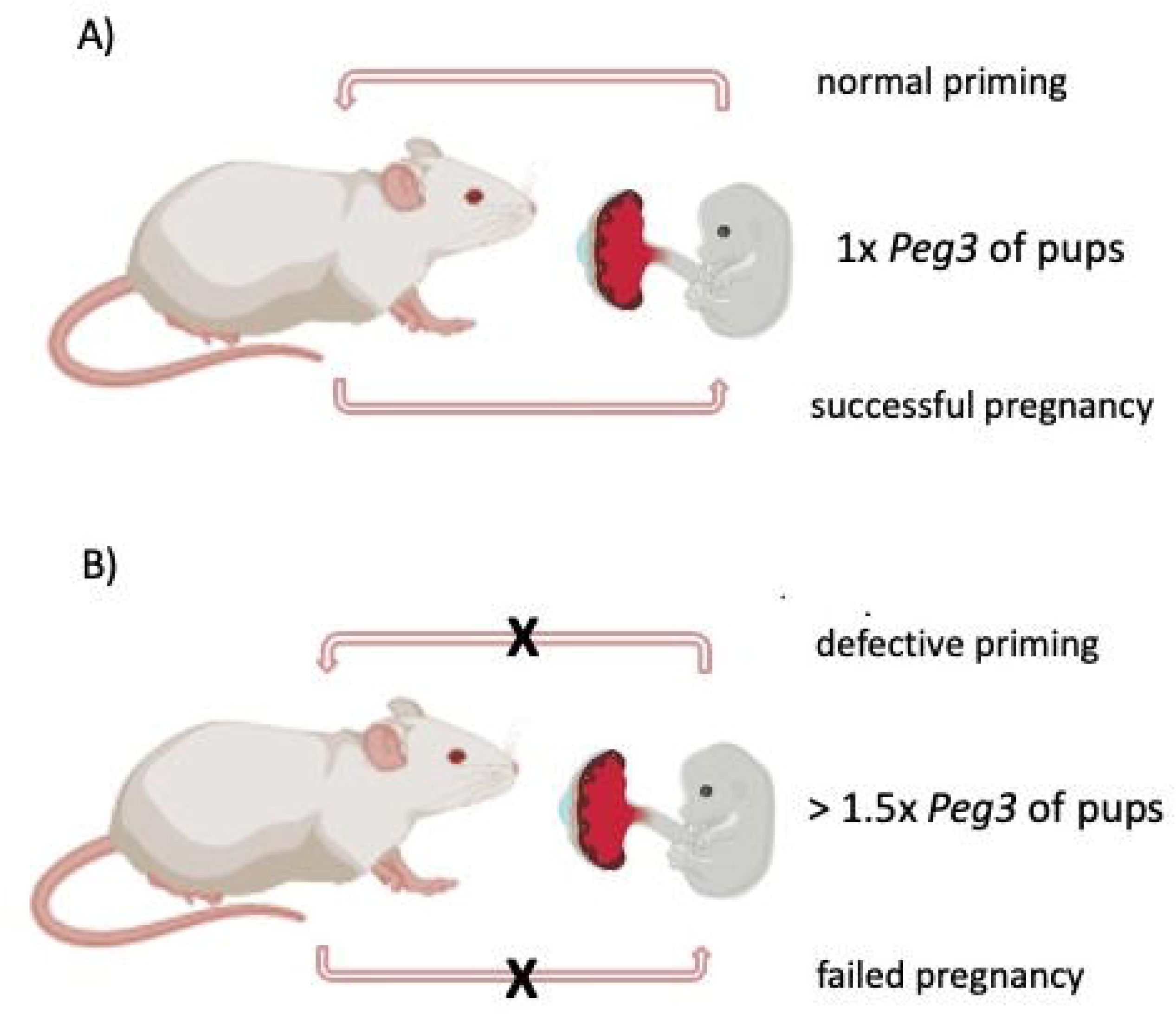
Fetal influence on maternal reproductive physiology *via Peg3*. (**A**) In mammalian pregnancy, fetal signals transmitted through the placenta prime the mother for parturition, placentophagy, nest building, and nursing behaviors. A normal fetal *Peg3* dosage (1x) properly primes the mother and supports successful pregnancy. (**B**) An elevated fetal *Peg3* dosage (>1.5x) may disrupt this priming process, resulting in delayed parturition and incomplete placentophagy, finally causing neonatal lethality.

The present study also uncovered evidence that female embryos are more vulnerable than males to elevated *Peg3* dosage (**Figure 5**). Although this sex-specific lethality was unexpected, several previous studies are consistent with the idea that the biological functions of the *Peg3* domain may be sexually biased. For instance, expression levels of *Peg3*and *Zim1* are higher in males than females during embryogenesis [34]. Additionally, earlier studies using loss-of-function mutations in *Peg3* (0x dosage) revealed more severe phenotypic effects in males than in females [11,12]. These findings suggest that optimal *Peg3* expression levels differ between the sexes, implying that males and females may possess distinct tolerable ranges of *Peg3* dosage. Under this model, a severe reduction in *Peg3* expression would be more detrimental to males, whereas a severe increase—as observed in the U1 strain—would be more detrimental to females. This framework aligns with the current observation that female embryos with 2x *Peg3* dosage exhibit disproportionately high lethality. Although female-specific lethality was not initially anticipated, the results are consistent with the broader hypothesis that the *Peg3* imprinted domain functions in a sex-biased manner [34].

The U1 allele is predicted to exert its mutational effects through dosage changes across the entire domain rather than through *Peg3* alone (**Figure 1**). Maternal transmission of the U1 allele results in the 2x dosage of the paternally expressed genes (*Peg3, Usp29, Zfp264, APeg3*) and the concurrent 0x dosage of the maternally expressed genes (*Zim1, Zim2, Zim3*) [13]. Therefore, the results from the current study must be interpreted with caution, because the extent to which the dosage change of each individual gene contributes to the observed phenotypes remains unclear. It is also important to consider the unusual evolutionary pattern associated with this imprinted domain. Although the genomic organization is highly conserved across mammalian species, several genes in the domain have lost their open reading frames (ORFs) in a lineage-specific manner during evolution—including *Zim2*, *Zim3*, and *Zfp264* in rodents, and *Usp29* in primate and artiodactyl lineages [35-37]. Despite losing their ORFs, these genes maintain transcriptional activity, suggesting that they may now function as non-coding RNA genes within each lineage [10]. In contrast, *Peg3* is the only gene that has retained its ORF throughout mammalian evolution, consistent with the idea that *Peg3* performs an essential function for species survival. This notion is further supported by the fact that similar phenotypic outcomes are observed in two different types of mutant strains: those targeting the *Peg3* ORF and those targeting the entire domain, as exemplified by the KO2 strain, which deletes the Peg3-DMR. Among these shared phenotypes, reduced growth rates are the most prominent [11-15]. If the same principle applies to the U1 allele—which produces the 2x dosage of *Peg3* along with additional dosage changes in neighboring genes—then the major defects observed in the U1 strain, such as delayed parturition and defective placentophagy, may primarily result from the 2x *Peg3* dosage rather than from combined domain-wide effects. This interpretation is plausible given the known function of PEG3 as a DNA-binding protein that regulates a large number of target genes [31,32]. However, we cannot rule out the possibility that some of the observed phenotypes arise from dosage changes in other, less expected genes within the domain. Additional studies will be required to disentangle these contributions.

## Materials and methods

### Ethics Statement

All mouse experiments were performed in accordance with National Institutes of Health guidelines for the care and use of laboratory animals and were approved by the Louisiana State University Institutional Animal Care and Use Committee (IACUC), protocol #25-043.

### Mouse Breeding

Two previously characterized mutant mouse strains were used in this study: the U1 strain, which carries a 1-kb deletion of the oocyte-specific alternative promoter U1, and the KO2 strain, which carries a 4-kb deletion of the Peg3-DMR [12,13]. All mice were housed in the Division of Laboratory Animal Medicine (DLAM) at Louisiana State University under a 12:12 h light–dark cycle at a constant temperature of 70°F and 50% humidity. Animals were provided ad libitum access to water and Rodent Diet 5001. Breeding cages were set up by placing two females with one male immediately before the onset of the dark cycle (4–6 PM). The following morning (9–11 AM), females were examined for the presence of vaginal plugs. Females with plugs were designated as being at gestational day 0.5. Pregnant females were transferred to individual cages at gestational day 15.5 and monitored daily to record gestation length and to document reproductive phenotypes, including placentophagy and nest-building behavior. Timed-mating experiments were set up in the same manner. Pregnant females were euthanized between gestational days 12.5 and 17.5 to collect embryos at various developmental stages.

For genotyping, genomic DNA was isolated from ear biopsies following overnight incubation at 55°C in lysis buffer (0.1 M Tris-Cl, pH 8.8, 5 mM EDTA, pH 8.0, 0.2% SDS, 0.2 M NaCl, 20 μg/ml Proteinase K). PCR genotyping was performed using the following primers: for the U1 allele, P1 (5’-TAGCAAGGGAGAGGGCCTAG-3’), P2 (5’-GGAAGCCTCCATCCGTTTGT-3’), and P3 (5’-AGCACAGCTAGAAATACACAGA-3’); for the KO2 allele, Primer A (5’-TGACAAGTGGGCTTGCTGCAG-3’), Primer B (5’-GGATGTAAGATGGAGGCACTGT-3’), and Primer D (5’-AGGGGAGAACAGACTACAGA-3’); for sex determination, mSry-F (5’-GTCCCGTGGTGAGAGGCACAAG-3’) and mSry-R (5’-GCAGCTCTACTCCAGTCTTGCC-3’).

### DNA Methylation Analysis

Genomic DNA isolated from mouse tissues was subjected to bisulfite conversion using the EZ DNA Methylation Kit (Zymo Research), following the manufacturer’s instructions [38]. Bisulfite-converted DNA was analyzed by COBRA (Combined Bisulfite and Restriction Analysis) as described previously [20]. Briefly, the converted DNA was amplified by PCR using primers targeting a 345-bp region of the Peg3-DMR (chr7: 6,729,539–6,729,883; mm10): Peg3Met15.2 (5’-TAGGTAGTTAATTAGGATAAGTTTGTGTAG-3’) and Peg3Met16.1 (5’-CCTATTACAAAACCACCACAATAAACATCA-3’). PCR products were digested with *Hph*I, which recognizes the sequence GGTGA. Unmethylated genomic DNA converts the original GGCGA site to GGTGA after bisulfite treatment and is therefore digestible. In contrast, methylated CpG sites remain as GGCGA and cannot be digested. Digested products were separated by electrophoresis on a 2% agarose gel: digested products represent the unmethylated allele, and undigested products represent the methylated allele of the Peg3-DMR.

## Acknowledgements

I thank Drs. Pini Perera and Corey Bretz for their valuable insights and constructive feedback on this manuscript. I also acknowledge Dr. Corey Bretz for his earlier contributions to the characterization of the U1 mutant strain.

